# Prolonged dexamethasone exposure enhances zebrafish lateral-line regeneration but disrupts mitochondrial homeostasis and hair cell function

**DOI:** 10.1101/2022.04.19.488823

**Authors:** Allison L. Saettele, Hiu-tung C Wong, Katie S Kindt, Mark E Warchol, Lavinia Sheets

## Abstract

The synthetic glucocorticoid dexamethasone is commonly used to treat inner ear disorders. Previous work in larval zebrafish has shown that dexamethasone treatment enhances hair cell regeneration, yet dexamethasone has also been shown to inhibit regeneration of peripheral nerves after lesion. We therefore used the zebrafish model to determine the impact of dexamethasone treatment on lateral-line hair cells and primary afferents. To explore dexamethasone in the context of regeneration, we used copper sulfate (CuSO4) to induce hair cell loss and retraction of nerve terminals, and then allowed animals to recover in dexamethasone for 48 hours. Consistent with previously work, we observed significantly more regenerated hair cells in dexamethasone-treated larvae. Importantly, we found that the afferent processes beneath neuromasts also regenerated in the presence of dexamethasone and formed an appropriate number of synapses, indicating that innervation of hair cells was not inhibited by dexamethasone. In addition to regeneration, we also explored the effects of prolonged dexamethasone exposure on lateral-line homeostasis and function. Following dexamethasone treatment, we observed hyperpolarized mitochondrial membrane potentials (ΔΨm) in neuromast hair cells and supporting cells. Hair cells exposed to dexamethasone were also more vulnerable to neomycin-induced cell death. In response to a fluid-jet delivered saturating stimulus, calcium influx through hair cell mechanotransduction channels was significantly reduced, yet presynaptic calcium influx was unchanged. Cumulatively, these observations indicate that dexamethasone enhances hair cell regeneration in lateral-line neuromasts, yet also disrupts mitochondrial homeostasis, making hair cells more vulnerable to ototoxic insults and possibly impacting hair cell function.

## Introduction

Glucocorticoids are the major hormones of the stress response pathway. Through interactions with ubiquitously expressed receptors, glucocorticoids are involved in regulating many physiological processes, including circadian cycle, tissue homeostasis, metabolism, and inflammation (Vettorazzi et al., 2021). Synthetic glucocorticoids, such as dexamethasone, are commonly used to treat inner ear disorders, such as sudden hearing loss and Menier’s disease, and have been proposed as an intervention to protect against cisplatin ototoxicity (Waissbluth et al., 2013). Achieving therapeutic levels of synthetic glucocorticoids in the inner ear with systemic oral delivery of the drug requires high-dose therapy associated with adverse side effects (Egli Gallo et al., 2013). Furthermore, systemic delivery of glucocorticoids to prevent cisplatin ototoxicity may render some types of malignant cells insensitive to cisplatin chemotherapy (Ge et al., 2012). Consequently, local intratympanic delivery directly into the ear has been tested as a potential method for the prevention and treatment of inner ear disease (Marshak et al., 2014; Patel, 2017). One limitation to this approach is that dexamethasone is rapidly cleared from the middle ear (Salt et al., 2011). Thus, strategies to provide sustained delivery of dexamethasone to the inner ear are an active area of research (Fernandez et al., 2016; Liao et al., 2021). While retention of synthetic glucocorticoids in the ear may provide greater therapeutic benefits, the effects of prolonged dexamethasone exposure on hair cell organ homeostasis and function are not well defined.

The zebrafish model has been used in numerous studies evaluating the effects of dexamethasone exposure on biological processes, including wound repair and complex tissue regeneration. In a previous study, dexamethasone was identified in a large-scale screen as an enhancer of hair cell regeneration after neomycin damage to zebrafish lateral-line organs, as well as inducing the addition of hair cells to lateral-line neuromasts in the absence of damage (Namdaran et al., 2012). By contrast, treatment with dexamethasone has been shown to inhibit peripheral nerve and spinal motor nerve regeneration in zebrafish (Bremer et al., 2017; Ohnmacht et al., 2016). Cumulatively, these data suggest that, while dexamethasone potentiates the generation of lateral-line hair cells, reinnervation of dexamethasone-induced hair cells may be impaired.

In this study we sought to address two questions: 1) does dexamethasone treatment enhance or inhibit afferent innervation and synapse formation in regenerating lateral-line organs, and 2) does prolonged dexamethasone exposure affect hair cell homeostasis and function? Our results reveal that dexamethasone treatment does not inhibit either afferent innervation or synapse formation in newly produced hair cells. However, dexamethasone treatment does increase mitochondrial activity in existing hair cells and reduces their evoked mechanosensory responses, suggesting that prolonged dexamethasone exposure may adversely affect hair cell function.

## Materials and Methods

### Ethics statement

Experimental procedures were performed with approval from the Washington University School of Medicine or NIH Institutional Animal Care and Use Committee, and in accordance with NIH guidelines for use of zebrafish.

### Zebrafish

Adult zebrafish were raised under standard conditions at 27-29°C in the Washington University or NIH Zebrafish Facility. Embryos were raised in incubators at 28°C in embryo media (EM; 15 mM NaCl, 0.5 mM KCl, 1 mM CaCl_2_, 1 mM MgSO_4_, 0.15 mM KH_2_PO_4_, 0.042 mM Na_2_HPO_4_, 0.714 mM NaHCO_3_) with a 14 h:10 h light:dark cycle. At 3-4 days post fertilization (dpf), larvae were anesthetized in 0.02% tricaine methanesulfonate (MS-222) and screened for transgenic fluorophores using a Leica MZ10 F stereomicroscope with fluorescence equipped with a GFP and a DSR filter set. After screening, 4 dpf, larvae were raised in 100-200 ml EM media in 250-ml plastic beakers and fed rotifers daily. Sex of the animal was not considered for this study because sex cannot be determined in larval zebrafish.

### Transgenic lines

The following transgenic lines were used in this study: nl1Tg *(TgBAC(neurod1:EGFP))*(Obholzer et al., 2008); w78Tg (*Tg(myo6b:GCaMP3)*) and w119Tg (*Tg(myo6b:mitoGCaMP3)*)(Esterberg et al., 2013), w200Tg *(Tg(mpeg1:yfp))* (Roca & Ramakrishnan, 2013), y229Gt *(Tg(tnks1bp1:EGFP))* (Behra et al., 2012) and idc1Tg *(Tg(myo6b:GCaMP6sCAAX))* (Sheets et al., 2017).

### Copper ablation of neuromasts and dexamethasone post-treatment

To ablate lateral-line neuromast hair cells and induce afferent neurite retraction, 5 dpf larvae were exposed to 3 μM CuSO_4_ (Sigma-Aldrich, Cat# 451677) diluted in EM in cell strainers (Corning; Cat# 431752) for 1 hour. This concentration and time course of CuSO_4_ exposure was verified as the optimal minimum dose to induce complete neuromast hair cell loss (mean hair cell number per neuromast: 11 ± 3 (control) vs 0 ± 1 (3 uM CuSO_4_), 14 fish per condition, 3 trials). After exposure, larvae were rinsed, and then allowed to swim freely during a 2-hour recovery in EM. Immediately following this 2-hour recovery, larvae were moved to 6-well plates (Falcon, Cat# 351146) containing 4 mL EM per well with either 0.1% DMSO (drug carrier alone) or 0.1% DMSO + 10 μM dexamethasone (Sigma-Aldrich, Cat# D1756) for 48 hours. The dexamethasone concentration used in this study (10 μM) was based on the concentration used in a previous study that identified dexamethasone as an enhancer of lateral-line hair cell regeneration (Namdaran et al., 2012). Larvae were maintained at 28°C throughout CuSO_4_ exposure and dexamethasone post-treatments.

### Neomycin ablation of neuromasts with dexamethasone pre-treatment

Macrophages are effector cells of the innate immune system, and in a previous study we observed robust recruitment of macrophages to neomycin-injured neuromasts (Warchol et al., 2021). To test whether dexamethasone treatment inhibited macrophage recruitment following hair cell injury, 5 dpf w200Tg *(Tg(mpeg1:yfp))* larvae were pre-treated with 0.1% DMSO or 1-50 μM dexamethasone in EM for 24 hours. Following pre-treatment, larvae were rinsed twice and placed in cell strainers, and then exposed to 100 μM neomycin (Sigma-Aldrich, Cat# N1876) for 30 minutes. Following neomycin exposure, fish were rinsed and moved to 6-well plates with 4 mL of EM to recover for 1 hour. Exposure and recovery times for neomycin treatment were chosen based on previous studies evaluating aminoglycoside toxicity in zebrafish lateral-line hair cells (Coffin et al., 2013; Owens et al., 2009).

To test whether dexamethasone pre-treatment sensitized hair cells to ototoxic insults, 5 dpf wild-type larvae were pre-treated with 0.1% DMSO or 10 μM for 48 hours, then exposed to 0-100 μM neomycin as described above.

### Whole-mount Immunohistochemistry

Free-swimming larvae were exposed to DAPI (1:2000, 4 minutes; D3571, Thermo Fisher) to label hair cell nuclei, then sedated on ice for 4 min. For GFP immunolabeling to label the afferent process or for Otoferlin immunolabeling to label hair cells, larvae were fixed (4% paraformaldehyde, 4% sucrose, 150 μM CaCl_2_, 0.1 M phosphate buffer) in 2 mL Eppendorf tubes overnight at 4-8°C. For synaptic labeling, larvae were fixed for 5 hours at 4-8°C.

Larvae fixed overnight were blocked in phosphate-buffered saline (PBS: 8 g NaCl, 0.2 g KCl, 0.24 g KH_2_PO_4_, 14.4 g Na_2_HPO_4_ per 1 L distilled H_2_O) with 5% normal horse serum (NHS; Sigma-Aldrich, Cat# H0146), 1% DMSO, 1% Triton X-100 (Sigma-Aldrich, Cat# T8787) for 2 hours at room temperature. Primary antibody exposure was performed in PBS with 2% NHS, 0.1% Triton-X100. Secondary antibody exposure was performed in PBS with 2% NHS. Following all antibody exposures, larvae were rinsed several times in PBS with 0.1% Tween-20 (Sigma-Aldrich, Cat# P9416).

For synaptic labeling, larvae were fixed for 5 hours, permeabilized in ice-cold acetone (−20°C, 5 min, Sigma-Aldrich, Cat# 179124), then blocked in PBS with 2% goat serum (Sigma-Aldrich, Cat# G9023), 1% bovine serum albumin (BSA; Sigma-Aldrich, Cat# A9647), and 1% DMSO for 2 hours. Primary and secondary antibody solutions and all post-antibody rinses were performed in the same PBS, BSA, and DMSO concentration mix used for blocking.

Primary antibodies:

- Mouse IgG2a α-CtBP (1:2000; Santa Cruz Biotech, Inc; Cat#: sc-55502)
- Mouse IgG1 α-MAGUK (K28/86; 1:500; NeuroMab, UC Davis; Cat# 75-029)
- Chicken α-GFP (1:500; Aves Labs, Inc; Cat# GFP-1020)
- Mouse IgG2a α-Otoferlin (1:500; Developmental Studies Hybridoma Bank; HCS-1)

Secondary antibodies (used at 1:1000):

- Goat α-chicken (Alexa 488) - neurites
- Goat α-mouse IgG1 (Alexa 488) – MAGUK
- Goat α-mouse IgG1 (Alexa 647) - MAGUK
- Goat α-mouse IgG2a (Alexa 647) - CtBP
- Goat α-mouse IgG2a (DyLight 550/Alexa546) – CtBP

All fish were rinsed in PBS prior to being mounted on glass slides in elvanol (13% w/v polyvinyl alcohol, 33% w/v glycerol, 1% w/v DABCO (1,4 diazobicylo[2,2,2] octane) in 0.2 M Tris, pH 8.5) and #1.5 cover slips.

### FM1-43 labeling

To evaluate hair cell mechanotransduction (Toro et al., 2015), 7dpf larvae were anesthetized in EM containing 0.01% Tricaine (EMT), exposed to 3 μM FM1-43 (N-(3-triethylammoniumpropyl)-4-(4-(dibutylamino)-styryl) pyridinium dibromide; ThermoFisher, Cat# T3163) for 20 seconds, rinsed in EMT, and then mounted on a FluoroDish (World Precision Instruments, Inc., Cat# FD3510) in low-melt agarose (Invitrogen, Cat# 16520) containing 0.01% Tricaine (LMAT). The agarose was allowed 2.5 minutes to solidify before covering in EMT and imaging.

### TMRE labeling

To measure mitochondrial ΔΨm, y229Gt *(Tg(tnks1bp1:EGFP))* transgenic larvae were exposed to 250 nM tetramethylrhodamine ethyl ester perchlorate (TMRE, Thermo Fisher, Cat# T669) diluted in EM for 25 minutes while wrapped in foil, then rinsed, allowed to equilibrate for 10-15 minutes, and mounted on a FluoroDish in LMAT. The agarose was allowed 2.5 minutes to solidify before covering in EMT and imaging.

### Confocal imaging

Most confocal images were acquired via an ORCA-Flash 4.0 V3 camera (Hamamatsu) using a Leica DM6 Fixed Stage microscope with an X-Light V2TP spinning disc confocal (60 micron pinholes) equipped with an LDI-7 laser diode illuminator (89 North). Image acquisition was controlled by MetaMorph software. *For fixed imaging:* z-stack images ~10-15 μm depth with a z-step of 0.2 μm were acquired using a 63x/1.4 N.A. oil immersion objective. *For live imaging*: larvae were anesthetized with 0.01% tricaine in EM, then mounted lateral-side up on a thin layer of 1% low-melt agarose in a tissue culture dish with a cover-glass bottom (FluoroDish; World Precision Instruments) and covered in EMT media. Z-stack images of neuromast L3-5 were acquired with a 63x/0.9 N.A. water immersion objective. Z-acquisition parameters with X-light spinning disc: z step of 1 μm, laser “20% power” for all wavelengths, 100 ms per frame for fixed imaging, 120 ms per frame for live imaging. Images in Fig. 5 were acquired using an LSM 700 laser scanning confocal microscope with a 63x 1.4 NA Plan-Apochromat oil-immersion objective (Carl Zeiss). Confocal z-stacks of ~15 μm depth were acquired with a z-step of 1 μm.

### Confocal image processing and analysis

Fiji (ImageJ software) was used to process and analyze most confocal images (Schneider et al., 2012). To analyze macrophage activation and inflammation, confocal images were reconstructed and analyzed using Volocity software (Quorum Technologies). Subsequent image processing for display within figures was performed using Illustrator software (Adobe).

Hair cell counts were quantified from hair cell specific DAPI labeled nuclei or from cells immunolabeled for Otoferlin (HCS-1; Fig. 5). To quantify hair cell synapses, raw images containing single immunolabel were converted to 8-bit, then subtracted for background using a 20-pixel (CtBP) or 10-pixel (MAGUK) rolling ball radius. Puncta were defined as regions of immunolabel with pixel intensity above a determined threshold: threshold for Ribeye label was calculated using the Isodata algorithm and threshold for MAGUK label was calculated using the Yen algorithm. Intact synapses were defined as CtBP immunolabel juxtaposed with MAGUK and quantified by scrolling through image z-stacks.

To measure neurite length, the ImageJ plugin Simple Neurite Tracer (Longair et al., 2011) was used on z-stack images of GFP-labeled afferent neurons. Following ImageJ’s Basic Instructions, which can be found in the Help drop-down menu of the Tracer plugin, neurites were traced by scrolling through the z-stack and creating paths. All neurite structures making up the basket-like processes that surround the neuromast were included in the paths, as well as the stalk at the base of the neuromast where innervating neurites emerged from the main afferent neuronal tract and any meandering projections coming off the dendritic basket. The Simple Neurite Tracer automatically detected the path set between two designated points and calculated a measured length in microns. Any branches emerging from and between paths were measured as separate paths. Once fully traced, all paths within the dendritic basket were then added together to get the total neurite length within the neuromast.

Macrophage density was quantified by scrolling through image z-stacks and counting the number of macrophages within 25 μm radius of a neuromast (using a circle inscribed on the neuromast). The number of macrophages contacting a neuromast was determined by scrolling through the x-y planes of each image stack (1 μm interval between x-y planes, 15 μm total depth) and counting macrophages that were in direct contact with Otoferlin-labeled hair cells. Finally, the number of macrophages that had internalized Otoferlin-immunolabeled material (hair cell debris) was counted and assumed to reflect the number of phagocytic events. For each metric, the recorded number reflected the activity of a single macrophage, i.e., a macrophage that made contacts with multiple hair cells and/or had internalized debris from several hair cells was still classified as a single ‘event.’

To measure the baseline fluorescence intensity of indicators across whole neuromasts, images containing single channels were background-subtracted using a rolling ball radius of the following sizes: images containing cytoGCaMp3 or mitoGCaMP3 (50-pixel), images containing FM1-43FX or TMRE label (100-pixel). Whole neuromasts were delineated based on hair cell specific DAPI label in maximum-intensity projections, and mean intensity of the indicator was measured. Measurements from each experimental trial were normalized to the control median value. To accommodate nested data and ensure statistical rigor, we preformed subsequent statistical analysis on average value of measurements from multiple neuromasts (L3, L4, L5) within an individual fish, such that each fish provided a single data point.

### Functional calcium imaging

GCaMP6s-based calcium imaging was used to examine evoked signals in zebrafish hair bundles and presynapses, as described previously (Lukasz & Kindt, 2018). Briefly, after 48-hour dexamethasone treatment, individual 7 dpf larvae were first anesthetized with EMT. Pins were used to restrain larvae onto a Sylgard-filled recording chamber. To suppress the movement, alpha-bungarotoxin (125 μM, Tocris) was injected into the cavity of the heart. Larvae were then immersed in extracellular imaging solution (in mM: 140 NaCl, 2 KCl, 2 CaCl_2_, 1 MgCl2 and 10 HEPES, pH 7.3, OSM 310 +/− 10) without tricaine. A fluid jet was used to mechanically stimulate hair bundles (L1-L4) with posterior fluid-flow for 500 ms.

To detect evoked calcium responses, images were acquired using a Bruker Swept-field confocal microscope (Bruker Inc), with a Nikon CFI Fluor 60x 1.0 NA water-immersion objective. A Rolera EM-C2 CCD camera (QImaging) was used to detect signals. To measure GCaMP6sCAAX signals, images were acquired with 2×2 binning using a 35 μm slit at 10 Hz in z-stacks of 5 planes every 2 μm or 0.5 μm for presynaptic and hair bundle measurements. High speed imaging along the z-axis was accomplished by using a piezoelectric motor (PICMA P-882.11–888.11 series, Physik Instrumente GmbH). Sequential evoked GCaMP6sCAAX acquisitions were separated by at least 5 min.

To quantify the magnitude of evoked GCaMP6s signals, images were processed in FIJI. Z-stacks were max-projected, and images at each timepoint were aligned using StackReg (Thevenaz et al., 1998). A 5 μm and 1.31 μm diameter circular ROI was drawn over each hair cell base and hair bundle respectively to make intensity measurements. To plot evoked changes in GCaMP6s intensity, the baseline (F_0_, signal during the 2 s pre-stimulus period) was used to plot the relative change in fluorescent signal from baseline (ΔF/F_0_). The magnitude of evoked calcium signals was determined using ΔF/F_0_ plots. The baseline GCaMP6s signal F_0_ was used to estimate the resting calcium levels in each hair bundle or presynaptic region. The response magnitudes or resting GCaMP6s levels for each neuromast were averaged to obtain a single point per neuromast.

### Statistical analysis

Statistical analyses were performed using Prism 9 (Graphpad Software Inc). Datasets were confirmed for normality using the D’Agostino-Pearson test—all the datasets in this study were normally distributed. Statistical significance between two groups was determined by an unpaired Student’s t-test if the two populations had the same variance, or an unpaired Welch’s unequal variance t-test if the two populations did not have the same variance. Statistical significance between multiple groups with normal distributions was determined by one-way ANOVA and appropriate post-hoc tests. For datasets dependent on multiple independent variables, statistical significance was determined using two-way ANOVA and appropriate post-hoc tests.

## Results

### Dexamethasone enhances regeneration and reinnervation of neuromast hair cells following CuSO_4_ lesion

In two separate small molecule screens using zebrafish larvae, dexamethasone has been identified as an enhancer of hair cell regeneration in lateral-line neuromasts and an inhibitor of peripheral fin nerve regeneration (Bremer et al., 2017; Namdaran et al., 2012). Dexamethasone has also been reported to enhance axon regeneration in rats, following injury (Feng & Yuan, 2015). We therefore addressed whether dexamethasone influences afferent reinnervation and hair cell synapse regeneration in zebrafish lateral-line neuromasts. To induce injury, we exposed animals to copper sulfate (CuSO_4_), which induces loss of lateral-line hair cells and considerable retraction of afferent nerve terminals (Hardy et al., 2021; Mackenzie & Raible, 2012; Olivari et al., 2008). Five-day old zebrafish larvae were treated in either 3 μM CuSO_4_ in embryo media (EM) or EM alone for 1 hour. After CuSO_4_ exposure, the animals were allowed to recover for 2 hours, and then transferred to EM containing either 10 μM dexamethasone or drug carrier alone (0.1% DMSO) for 48 hours (Fig. 1A). After 48 hours we assessed the extent of afferent reinnervation and hair cell and synapse regeneration. We focused on posterior lateral-line neuromasts L3-L5 for analysis (Fig. 1B). To examine afferent reinnervation and hair cell regeneration we visualized afferent nerve fibers labeled with GFP as well as hair cell nuclei labeled with DAPI (Fig. 1C). In agreement with previous observations of lateral-line organs injured with neomycin, we observed a significantly greater number of regenerated hair cells per neuromast in larvae exposed to dexamethasone during regeneration compared to controls (Fig. 2 C-E; orange points, ****adjusted P<0.0001; Tukey’s multiple comparisons test). Additionally, among fish that were not exposed to CuSO_4_ (unlesioned), significantly more hair cells per neuromast were observed in fish that were exposed to dexamethasone (Fig. 2 A,B,E; gray points). This observed increase in hair cell number per neuromast has been previously shown to be due to increased proliferation of hair cell precursors rather than prevention of hair cell turnover (Namdaran et al., 2012). These results therefore confirm that dexamethasone enhances hair cell regeneration and stimulates hair cell precursors to divide, even in the absence of injury.

**Figure 1.**
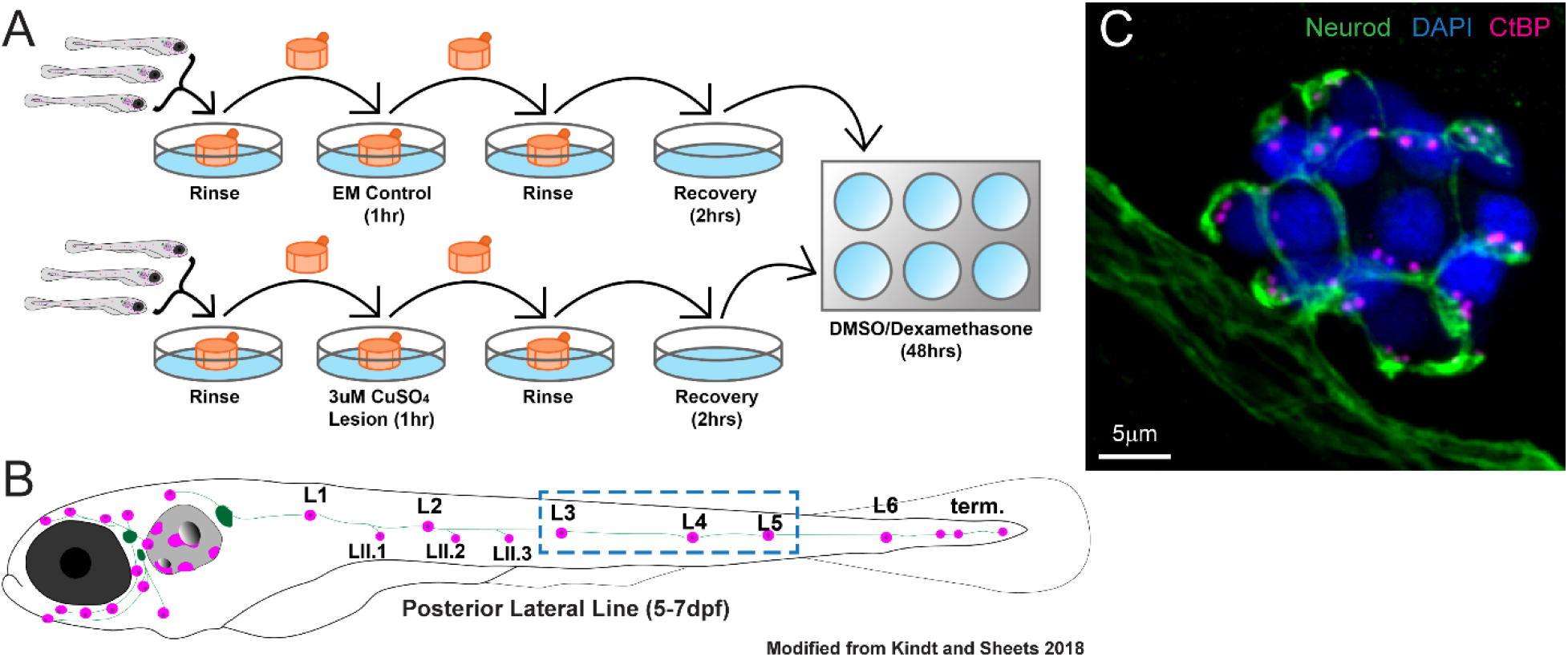
Workflow of CuSO_4_ lateral-line lesion and dexamethasone exposure. (A) Diagram of workflow for CuSO4 lesions and dexamethasone exposure. Larvae (5 dpf) were placed into cell strainers and moved between rinse and lesion steps, while being allowed to swim freely during recovery and dexamethasone exposure. (B) Diagram of a 5-7dpf zebrafish larvae showing the distribution of lateral line neuromasts (magenta) and innervating afferent nerves (green). This study focused on neuromasts L3, L4, and L5 (blue dashed-line box). (C) Representative maximum intensity projection of confocal z-stack image of neuromast L4 in a 7 dpf zebrafish larvae. Labels: afferent neurons labeled with GFP (green; tgBAC(neurod1:GFP), DAPI-labeled hair-cell nuclei (blue), and immunolabeled presynaptic ribbons (magenta; pan-CtBP).

**Figure 2.**
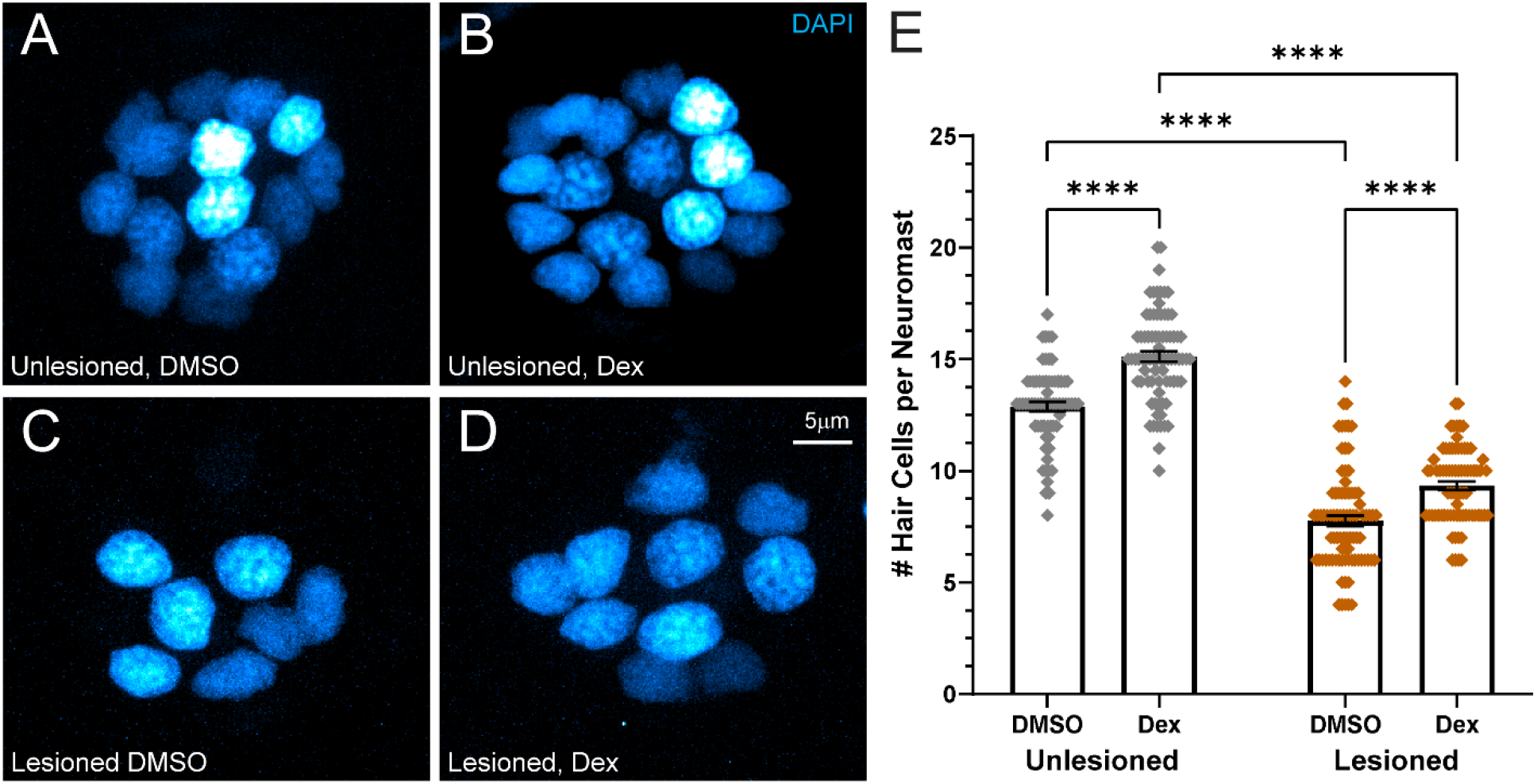
Dexamethasone increased hair cell number in intact and regenerating neuromasts. (A-D) Representative maximum intensity projection images of 7dpf neuromasts, with DAPI-labeled hair cell nuclei (blue). Larval lateral line neuromasts were either left intact (unlesioned) (A, B) or lesioned by treatment with 3μM CuSO_4_ for 1 hour (C, D). Larvae were then allowed to recover for 2hrs, then exposed for 48 hr to either 0.1% DMSO (carrier) (A, C) or dexamethasone (B, D). (E) Scatter plot graph of hair cell number per neuromast. Bars indicate mean and SEM. Each dot represents a single neuromast on an individual fish. Both intact (unlesioned) and regenerating neuromasts showed a significant increase in the number of hair cells per neuromast in dexamethasone-treated fish relative to DMSO controls (****adjusted P<0.0001). N= 5-10 fish per condition per trial; 7 experimental trials.

To determine whether dexamethasone treatment enhanced or inhibited afferent innervation of hair cells, we measured the length of innervating afferent processes in both unlesioned and lesioned neuromast after regeneration (Fig. 3 A-D). In neuromasts that had regenerated after CuSO_4_-induced lesions, we observed that the average length of afferent processes per neuromast was significantly greater in animals that recovered in dexamethasone compared to animals exposed to drug carrier alone (Fig 3 C-E (orange box plots); ****adjusted P<0.0001; Tukey’s multiple comparisons test). We reasoned that increased neurite length in dexamethasone-treated animals was related to the greater number of regenerated hair cells per neuromast. To verify this idea, neurite length was normalized relative to the number of hair cells in each neuromast. Following normalization, we observed no difference in relative neurite length per hair cell in animals recovered in dexamethasone vs. those in drug carrier alone (Fig 3F (orange box plots); adjusted P=0.5630). In unlesioned animals treated with dexamethasone, we observed no change in afferent neurite length per neuromast (Fig. 3 A,B,E (gray box plots); adjusted P=0.8215) and a modest reduction in the relative neurite length per hair cell that was not statistically significant (Fig 3F (gray box plots); adjusted P=0.2959). These data indicate that dexamethasone treatment enhanced the formation of afferent processes in regenerating neuromasts, perhaps to accommodate a greater number of regenerated hair cells. By contrast, in unlesioned neuromasts dexamethasone did not promote the addition of afferent neurites to accommodate additional hair cells.

**Figure 3.**
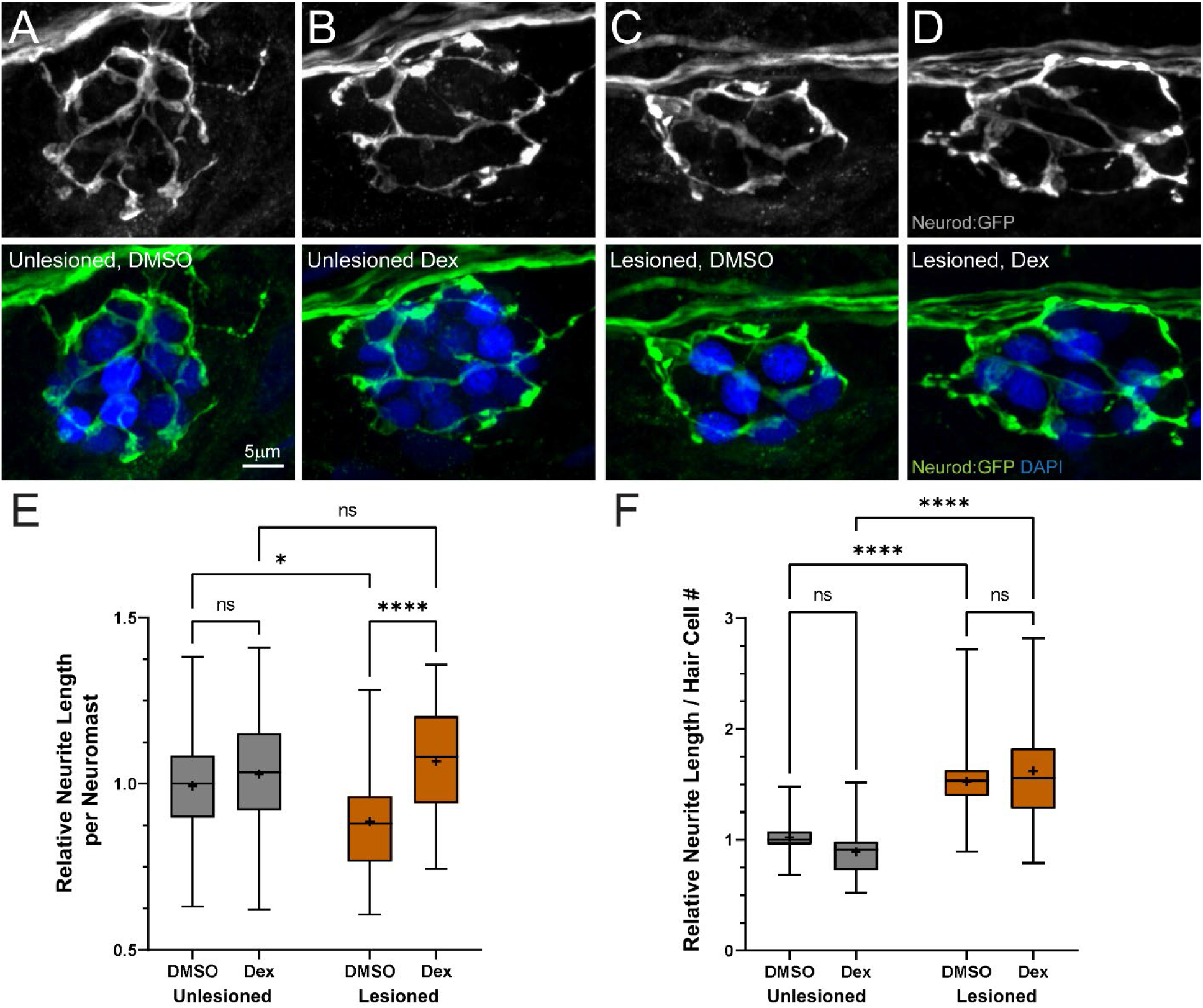
Dexamethasone enhanced afferent innervation of regenerating neuromasts. (A-D) Representative maximum intensity projection images of pLL neuromasts in 7dpf tgBAC(neurod1:GFP) larvae showing afferent neurons (green) and DAPI-labeled hair-cell nuclei (blue). Neuromasts were either unlesioned (A, B) or lesioned with CuSO_4_ (C, D), then allowed to regenerate for 48 hr while exposed to DMSO (carrier) (A, C) or dexamethasone (B, D). (E-F) Box plots of neurite length per neuromast (E) and neurite length per hair cell (F), with whiskers at min/max and mean values marked with “+”. Data for each experimental trial were normalized to the median value of unlesioned controls. Regenerating neuromasts (orange plots) in dexamethasone-treated fish had significantly longer neurites per neuromast (E) (****adjusted P<0.0001), however the difference in length in dexamethasone-treated fish was not significant relative to the total number of hair cells/neuromast, when compared to DMSO controls (F). N=4-7 fish per condition per trial; 6 experimental trials.

In addition to quantifying hair cell numbers and neurite length, we also quantified the number of intact synapses per hair cell (Fig. 4 A-D; defined as presynaptic ribbons adjacent to postsynaptic densities). We observed no difference in synapse counts between dexamethasone-treated and 0.1% DMSO controls when comparing unlesioned neuromasts or when comparing CuSO_4_ lesioned neuromasts following regeneration (Fig. 4 E). Notably, we did observe significantly more intact synapses per hair cell in all regenerating neuromasts relative to unlesioned neuromasts (Fig 4 E; adjusted ***P=0.0003 (DMSO), * P=0.0202 (Dex); Tukey’s multiple comparisons test). Similar observations have been reported in regenerating neuromast hair cells (Hardy et al., 2021; Suli et al., 2016) and may reflect ongoing synaptic refinement akin to that observed in developing cochlear hair cells (Michanski et al., 2019). Overall, these results suggest that dexamethasone treatment enhances afferent neurite outgrowth in regenerating neuromasts and does not impact synapse formation in newly formed hair cells in regenerating or unlesioned neuromasts.

**Figure 4.**
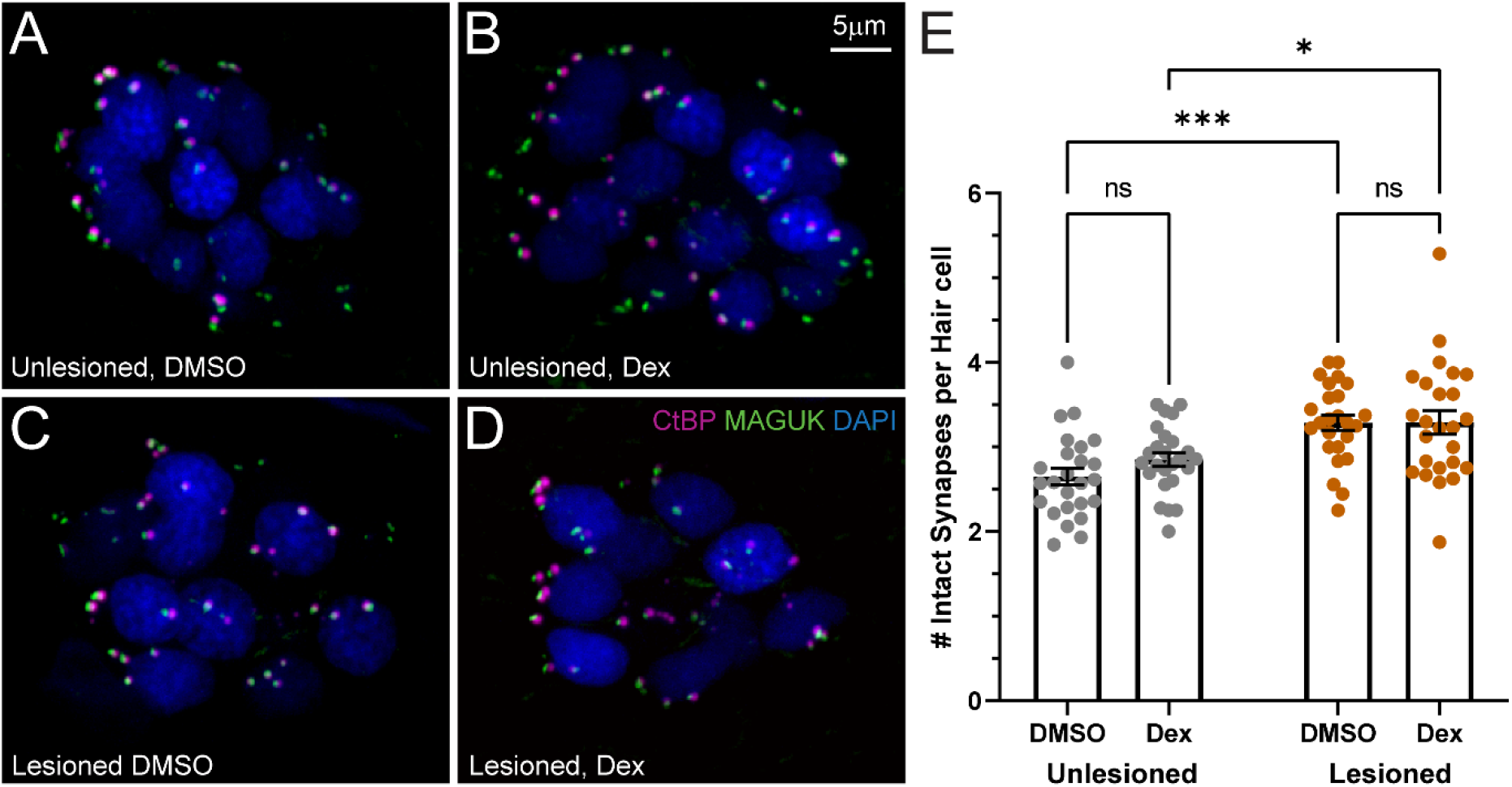
Dexamethasone exposure did not alter the number of afferent synapses in neuromast hair cells. (A-D) Representative maximum intensity projection images of 7dpf pLL neuromasts, with hair cell presynaptic ribbons labeled via CtBP antibody (magenta), post-synaptic densities labeled via MAGUK antibody (green), and hair cell nuclei labeled with DAPI (blue). Neuromasts were either unlesioned (A, B) or lesioned with CuSO_4_ (C, D), then allowed to regenerate for 48 hr while exposed to DMSO (carrier) (A, C) or dexamethasone (B, D). (E) Scatter plots with mean and SEM bars showing no significant difference in the number of intact synapses in dexamethasone-treated fish, compared to DMSO controls for both lesioned and intact neuromasts. Each dot represents a single neuromast on an individual fish.Notably, a significantly greater number of intact synapses per hair cell were observed in regenerating neuromasts relative to neuromasts that were not lesioned (adjusted ***P=0.0003 (DMSO), * P=0.0202 (Dex)). N=5 fish per condition per trial; 4 experimental trials.

### Pretreatment with dexamethasone does not inhibit macrophage response to hair cell injury

Dexamethasone inhibits inflammation and macrophage activation, and various studies indicate dexamethasone’s anti-inflammatory action can influence nerve regeneration (Valledor & Ricote, 2004). Previous work has shown that macrophages are recruited to lateral-line organs after injury induced by the ototoxin neomycin (Warchol et al., 2021). We therefore assessed whether dexamethasone inhibited macrophage entry and phagocytosis after application of the ototoxin neomycin. We quantified macrophage entry and phagocytosis in response to neomycin treatment after pretreating larvae for 24 hr with 1, 10 or 50 μM dexamethasone. Larvae that were not exposed to either neomycin and/or dexamethasone served as controls (Fig 5 A). All experiments followed the same protocol that we had previously used to characterize macrophage recruitment and phagocytosis after neomycin toxicity (Warchol et al., 2021). Dexamethasone treatment did not induce any significant changes to macrophage contacts in the absence of neomycin (Fig 5 B) or to the subsequent macrophage response following neomycin lesion (Fig. 5 C-F). Quantification revealed that the number of macrophage-hair cell contacts (Fig. 5 G) and number of phagocytotic events (Fig. 5 H) was similar in all neomycin treatment groups. These observations indicate that pre- and co-treatment with 10 μM dexamethasone does not suppress macrophage activation in response to neuromast injury.

**Figure 5.**
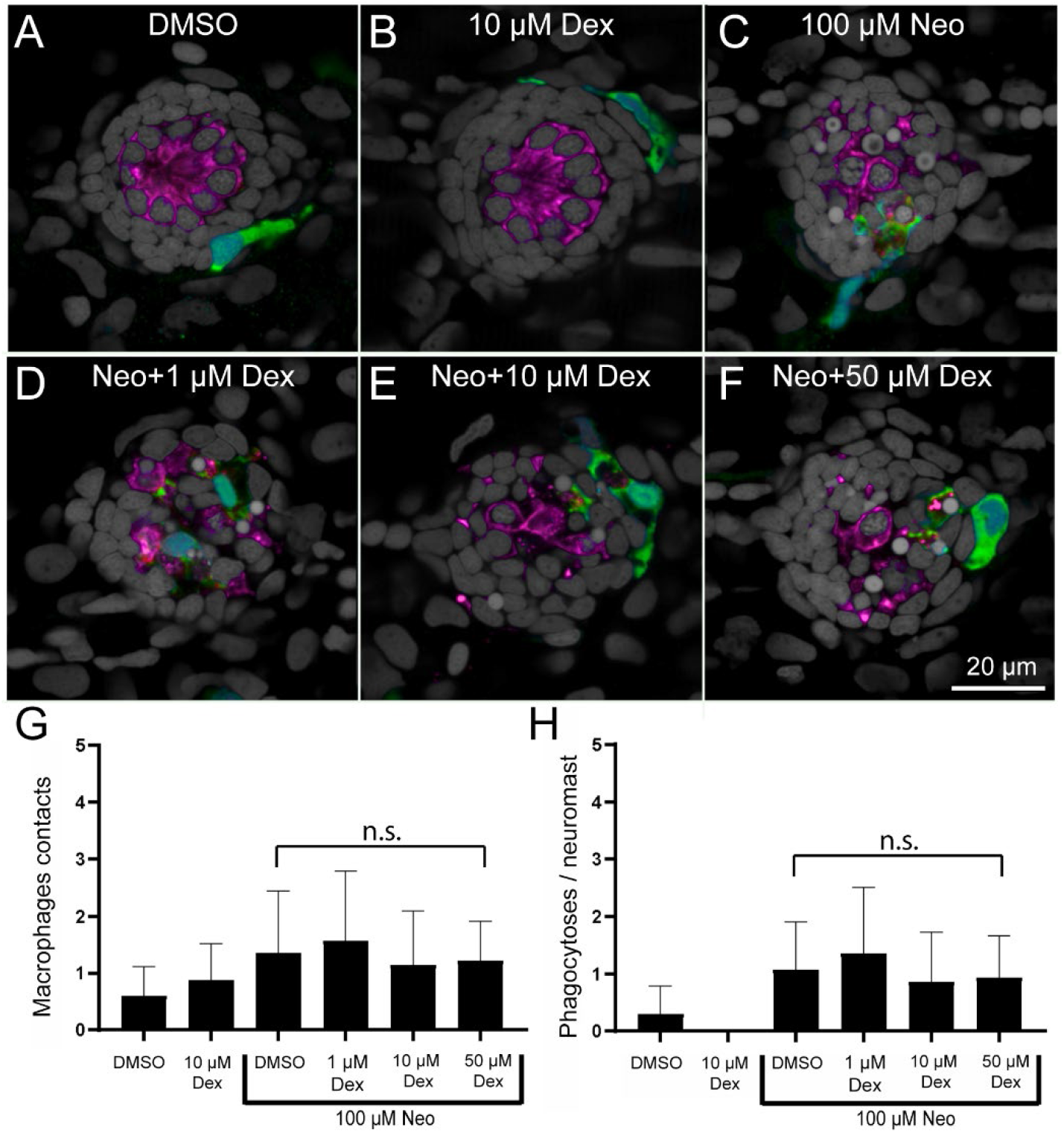
Dexamethasone did not inhibit macrophage response to neuromast injury. (A-F) Representative confocal images of control (A,B) and neomycin injured neuromasts (C-F). Larvae (6 dpf) were treated for 24 h with dexamethasone, then exposed for 30 min to neomycin. All specimens were fixed and processed for labeling of macrophages (green, YFP), hair cells (magenta, Otoferlin) and nuclei (gray, DAPI). Normal localization of macrophages was observed in control fish (A,B) and no inhibition was observed in neuromasts treated with neomycin (C-F). (G-H) Dexamethasone pretreatment did not affect macrophage contacts with dying hair cells (G, P>0.9999, Dunn’s multiple comparison test) or the number of phagocytotic events (H, P>0.9999, Dunn’s multiple comparison test). Bars indicate SD. N=15 fish/group; 2 experimental trials.

### Hair cell FM1-43 uptake and calcium homeostasis is unaffected by dexamethasone exposure

Numerous studies have indicated that dexamethasone can induce mitochondrial dysfunction in various tissues, including adipocytes (Luan et al., 2019), skeletal muscle (Liu et al., 2016), and neurons (Du et al., 2009). We previously reported reduced hair cell uptake of the cationic dye FM1-43 in zebrafish mutants with impaired hair cell mitochondrial function *(mpv17*^-/-^),indicating that these fish possessed reduced driving force for cations through mechanotransduction channels, and elevated hair cell mitochondrial calcium (Holmgren & Sheets, 2021). We therefore evaluated whether 48-hour dexamethasone exposure affected hair cell uptake of FM1-43 and calcium homeostasis. We observed no differences in FM1-43 uptake in unlesioned neuromasts that were treated in dexamethasone vs. those incubated in DMSO carrier (Fig. 6 A,B,E (gray circles)). In regenerating neuromasts, dexamethasone treatment appeared to cause a modest increase in FM1-43 uptake, but this trend did not reach statistical significance (Fig. 6 C-E (orange circles); adjusted P=0.1786; Tukey’s multiple comparisons test). Notably, hair cells in regenerating DMSO-treated neuromasts (Fig. 6 C) displayed significantly reduced FM1-43 uptake relative to unlesioned DMSO-treated neuromasts (Fig. 6 A,E; adjusted *P=0.0226), indicating reduced mechanotransduction in regenerated control neuromast hair cells. By contrast, there was no significant difference in FM1-43 uptake in regenerating vs. unlesioned dexamethasone-treated neuromast hair cells (Fig. 6 B,D,E; adjusted P=0.2747), suggesting that dexamethasone treatment during regeneration may enhance hair cell regeneration and may also slightly impact the maturation of mechanotransduction.

**Figure 6.**
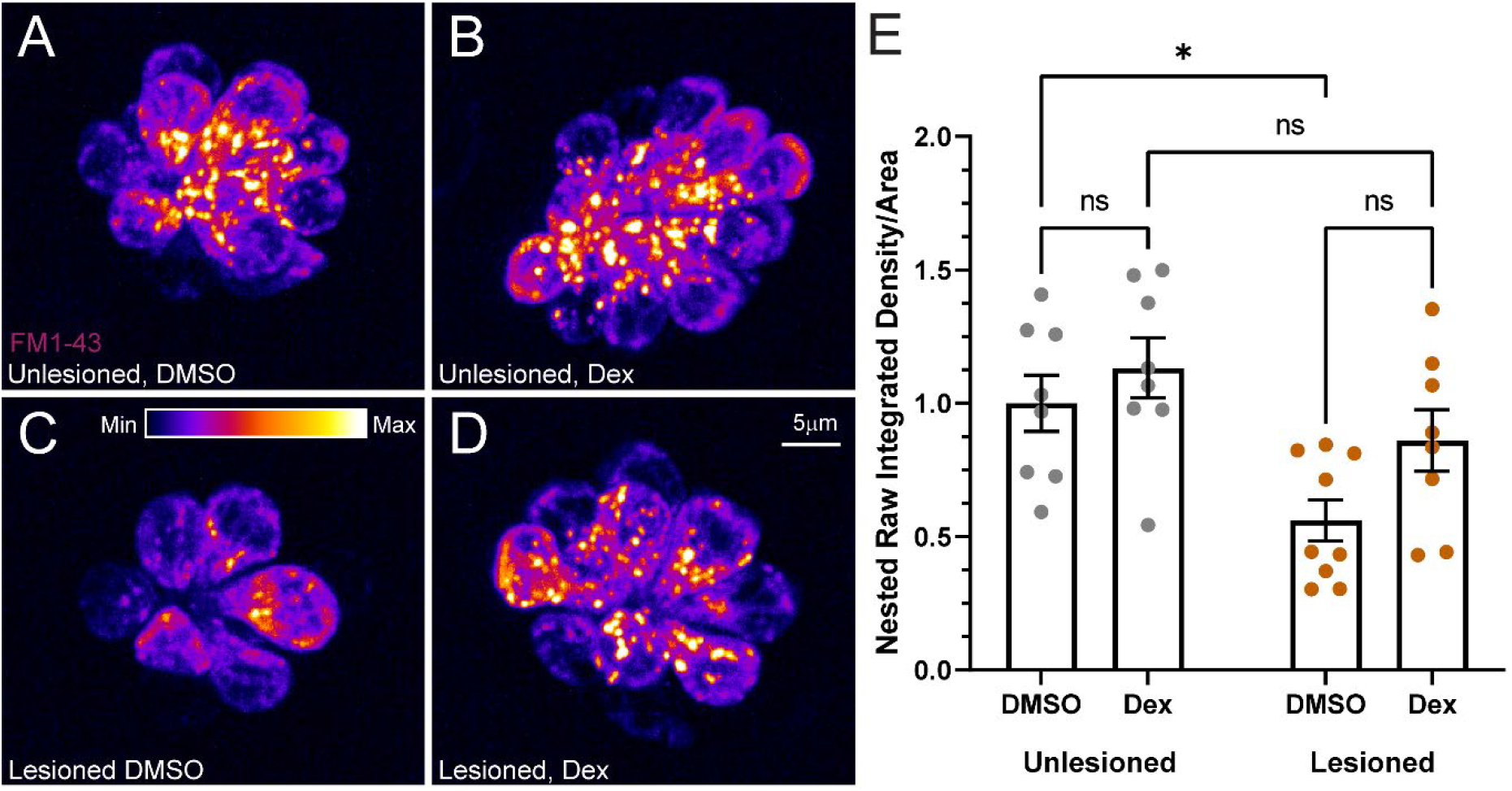
FM1-43 uptake was not significantly affected by dexamethasone exposure. (A-D) Representative maximum intensity projection images of pLL neuromasts in live 7dpf larvae immediately after brief (20 seconds) FM1-43 exposure. Neuromasts were either left intact (A, B) or lesioned with CuSO_4_ (C, D), then allowed to regenerate for 48 hr while exposed to DMSO (carrier) (A, C) or dexamethasone (B, D) prior to FM1-43 exposure. The LUT bar (C) shows the range of intensity values, starting at 0 (min) to 400 (max). (E) Relative average FM1-43 intensity per neuromast. Scatter plots show raw integrated density (i.e. sum of the pixel values)/area values with SEM bars. Each dot represents the average values of neuromasts L3-L5 for one fish. Values were normalized for each experiment using the inverse median of the unlesioned control values. The data show no significant differences in intensity for dexamethasone-treated fish, relative to DMSO controls, in both intact and regenerating neuromasts (adjusted *P=0.0226). N=2-3 fish per condition per trial; 4 experimental trials.

We next examined hair cell cytosolic and mitochondrial calcium homeostasis using the genetically encoded calcium indicator GCaMP3 (Esterberg et al., 2013; Esterberg et al., 2014). When comparing unlesioned neuromasts and CuSO_4_ lesioned neuromasts following regeneration, we observed average cytosolic calcium levels did not differ in neuromast hair cells (Fig. 7 A-E), while mitochondrial calcium levels appeared significantly higher in regenerating neuromast hair cells relative to unlesioned in both DMSO and dexamethasone treatment conditions (Fig. 7 F-J; adjusted ***P=0.0006 (DMSO), **P=0.0019 (Dex); Tukey’s multiple comparisons test). Notably, dexamethasone treatment had no effect on cytosolic or mitochondrial calcium levels relative to DMSO control in both unlesioned and regenerating neuromasts (Fig. 7 E, J). Cumulatively, these data suggest that prolonged dexamethasone exposure does not significantly impact hair cell mechanotransduction or affect calcium homeostasis in either intact or regenerating neuromasts.

**Figure 7.**
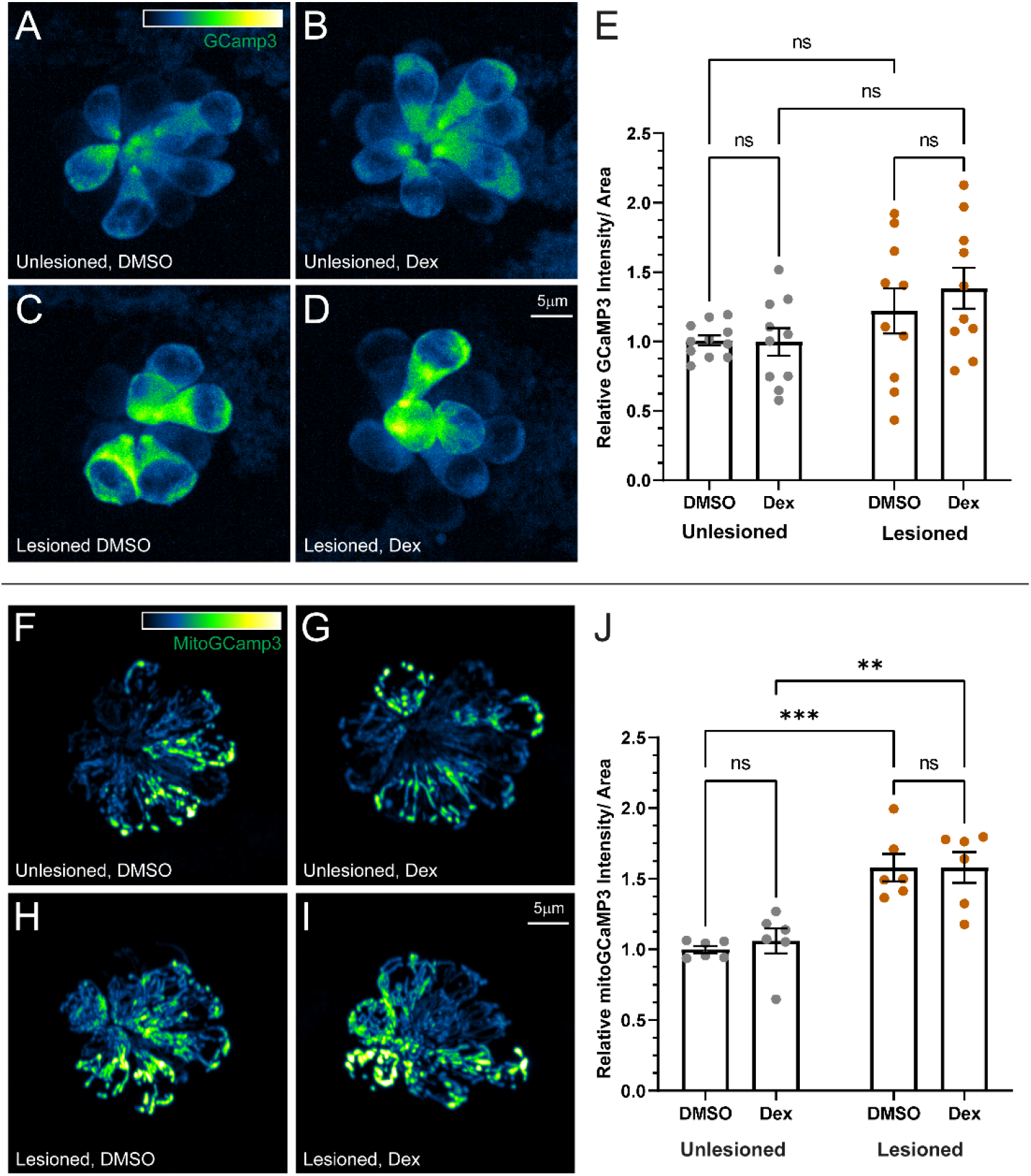
Dexamethasone did not affect hair cell cytosolic or mitochondrial calcium homeostasis. (A-D) Representative maximum intensity projection images of neuromasts in live 7dpf Tg(−6myo6b: GCaMP3) larvae. The LUT bar in (A) shows range of intensity values starting at 0 (min) to 2150 (max). (E) Hair cell GCaMP3 fluorescence intensity is not affected by dexamethasone exposure. Scatter plots of relative raw integrated intensity/area values with SEM bars. Each dot represents the average values of neuromasts L3-L5 for one fish. Data show no significant differences in intensity for dexamethasone-treated fish relative to DMSO controls in both intact and regenerating neuromasts. N=2-3 fish per condition per trial; 5 experimental trials (F-I) Representative images neuromasts in Tg(−6myo6b: mitoGCaMP3) larvae. LUT bar (F) shows range of intensity values starting at 0 (min) to 300 (max). (J) Hair cell mitoGCaMP fluorescence intensity is not affected by dexamethasone exposure but is elevated in regenerating neuromasts relative to intact neuromasts. Data show no significant differences in intensity for dexamethasone-treated fish relative to DMSO controls in both intact and regenerating neuromasts (adjusted ***P=0.0006 (DMSO), **P=0.0019 (Dex)). N=2-3 fish per condition per trial; 3 experimental trials.

### Prolonged dexamethasone exposure hyperpolarizes neuromast mitochondria and sensitizes hair cells to neomycin

To determine whether 48-hour dexamethasone exposure impacted mitochondrial function in lateral-line neuromasts, we examined mitochondrial membrane potentials (ΔΨm) in unlesioned neuromast hair cells and supporting cells using tetramethylrhodamine methyl ester (TMRE)—a cell-permeant cationic dye used to assess mitochondrial function (Alassaf et al., 2019; Esterberg et al., 2016). Dexamethasone-exposed neuromasts had significantly higher TMRE mean fluorescence intensity relative to controls (Fig. 8 A,B,E; **** P<0.0001; Welch’s t-test), indicating that mitochondria in dexamethasone-exposed neuromast cells are more hyperpolarized. Elevated TMRE fluorescence was apparent not only in the mitochondria of hair cells (Fig 8 A’, B’, F; EGFP-negative cells), but also the surrounding supporting cells (Fig 8 A”, B”, G; EGFP-positive cells), suggesting prolonged dexamethasone exposure disrupts normal mitochondrial function in both hair cells and supporting cells. Given that the hair cells in dexamethasone-treated larvae had more negatively charged ΔΨm, we predicted that these hair cells would be more vulnerable to neomycin-induced hair cell death (Alassaf et al., 2019; Holmgren & Sheets, 2021). We therefore compared neomycin-induced hair cell lesion in fish pre-treated with dexamethasone or DMSO carrier alone (Fig 9 A-J). When treated with higher doses of neomycin (50-100 μM), we observed significantly more lateral-line hair cell loss in fish pre-exposed for 48 hours to dexamethasone, relative to controls (Fig 9 K; adjusted P= 0.1407 (30 μM), *P=0.0353 (50 μM), ****P<0.0001 (100 μM); Šídák’s multiple comparisons test). Thus, while prolonged dexamethasone does not appear to impact cytosolic or mitochondrial calcium homeostasis, it does alter neuromast mitochondrial function and increases hair cell sensitivity to neomycin-induced death.

**Figure 8.**
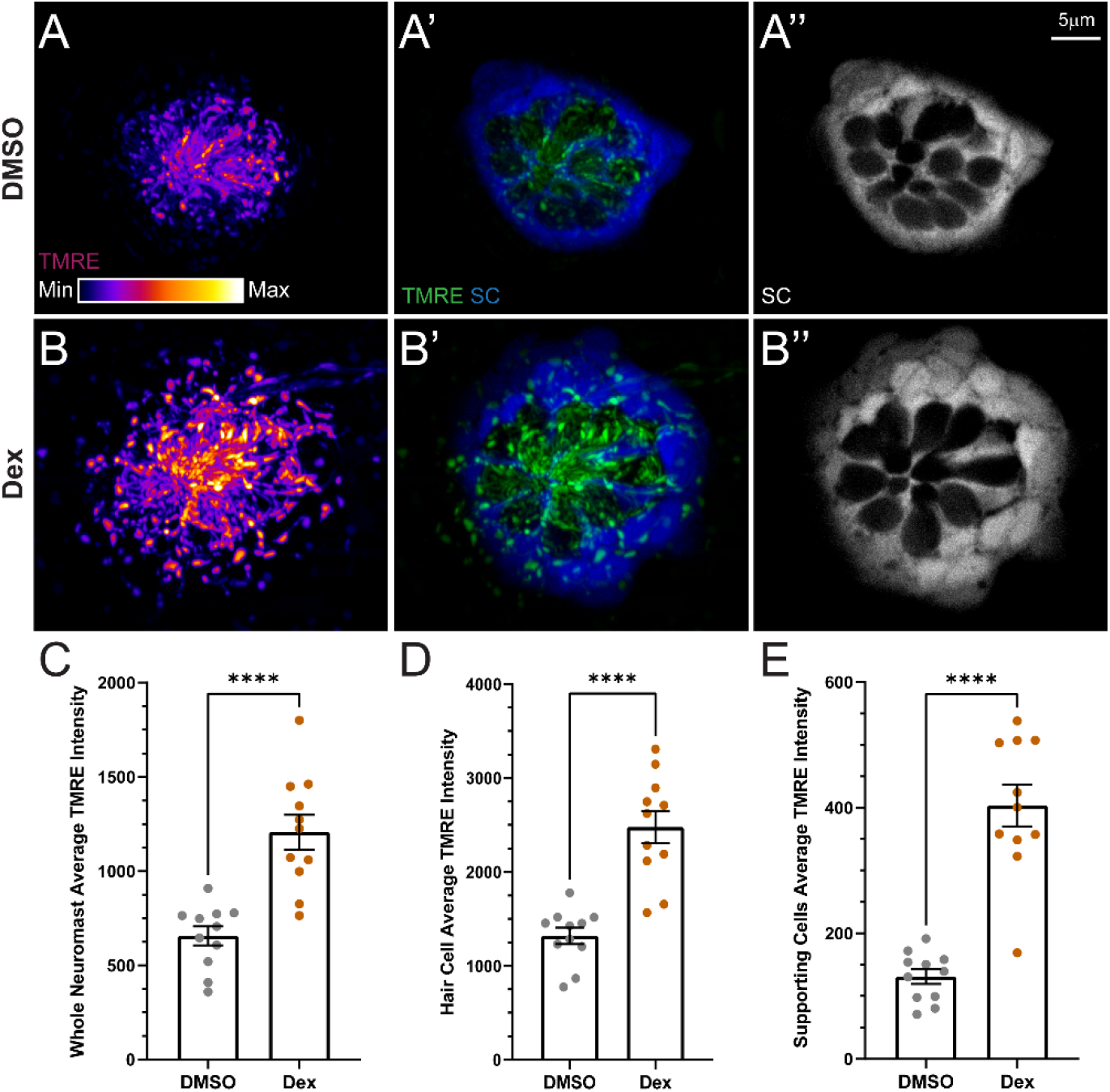
Dexamethasone exposure elevated mitochondrial activity in hair cells and supporting cells. (A-B) Confocal images of neuromasts in 7dpf fish following TMRE exposure. Fish were treated with DMSO (A) or dexamethasone (B) for 48 hours prior to imaging. TMRE (A, A’, B, B’) is shown as a max intensity projection and supporting cell (SC) images (A’, A’’, B’, B’’) are z-slices from the midlevel of the neuromast. Negative spaces inside GFP-labeled supporting cells indicate neuromast hair cells. LUT bar (A) shows range of intensity values starting at 0 (min) to 11800 (max). (E-G) Scatter plots of average TMRE fluorescence intensity with SEM bars. Dexamethasone-treated fish had significantly hyperpolarized mitochondria in both neuromast hair cells and supporting cells (adjusted **** P<0.0001). N=1-4 fish per condition per trial; 5 experimental trials.

**Figure 9.**
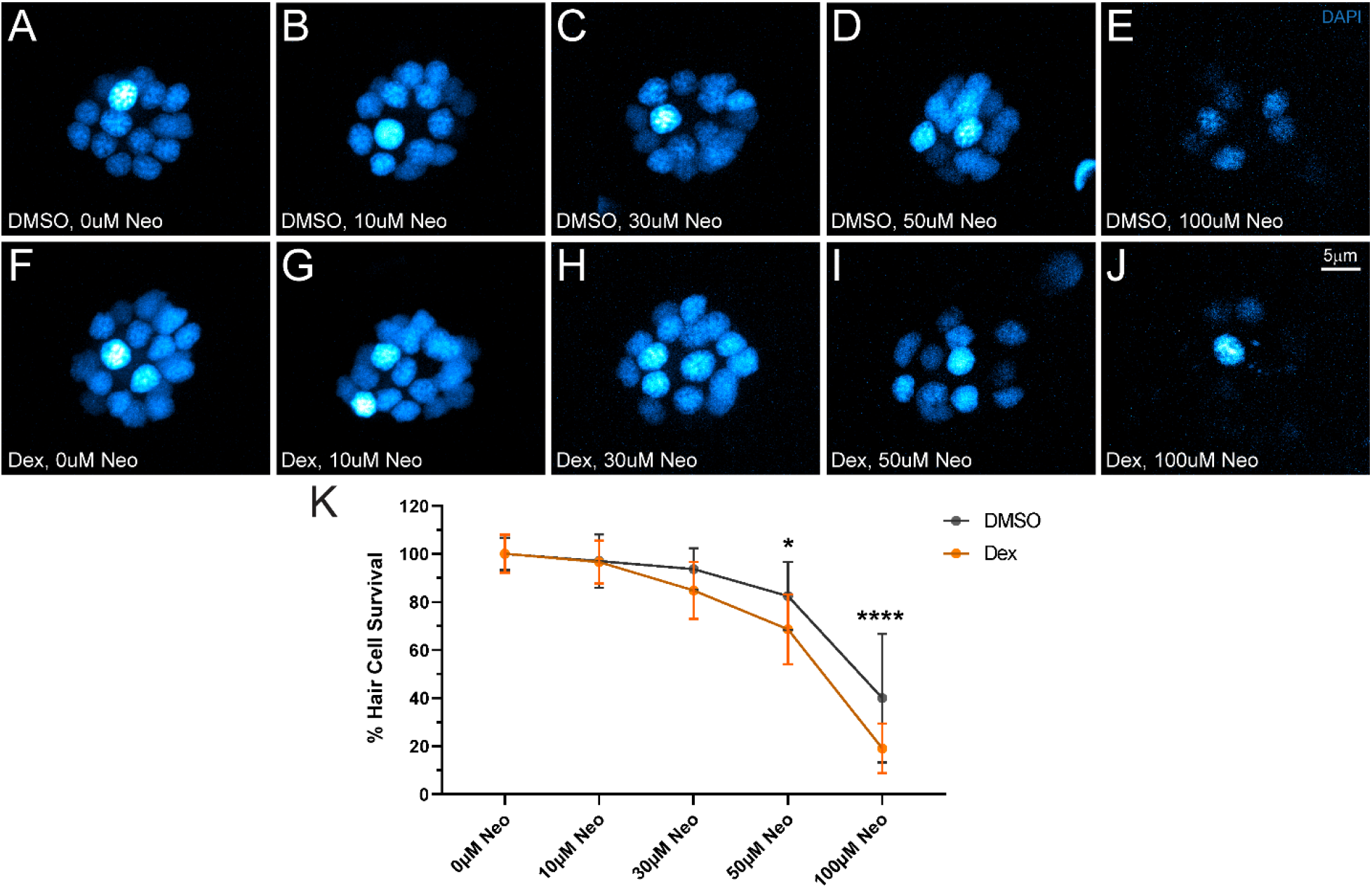
Dexamethasone exposure increased hair cell sensitivity to neomycin. (A-J) Representative maximum intensity images of DAPI-labeled neuromast hair cell nuclei (blue). Larvae (7dpf) were exposed for 48 hr to DMSO (carrier) alone (A-E) or dexamethasone (F-J), then exposed for 30min or 10μM (B, G), 30μM (C, H), 50μM (D, I), or 100μM (E, J) neomycin. EM buffer alone was used as a control (A,F). (K) Dose response curve with mean and SEM bars shows the number of surviving hair cells per neuromast as a percent of those present in unlesioned controls. Dexamethasone-treated neuromast hair cells were more sensitive to neomycin induced cell death than controls (adjusted P= 0.1407 (30 μM), *P=0.0353 (50 μM), ****P<0.0001 (100 μM)). N=7 fish per condition per trial; 3 experimental trials.

### Prolonged dexamethasone exposure reduces evoked calcium influx through mechanosensory hair bundles

Hair cells respond to sensory stimuli via deflection of apical hair bundles and opening of mechanotransduction channels. Influx of potassium and calcium occurs through these channels which depolarizes the hair cell, leading to influx of calcium through basally-localized presynaptic voltage-gated calcium channels, and triggering release of synaptic vesicles (Corey & Hudspeth, 1979; Moser & Beutner, 2000; Zhang et al., 2018). To determine if prolonged dexamethasone exposure influenced hair cell responses to mechanical stimulation, we measured evoked calcium influx at hair bundles and hair cell synapses in response to bundle deflection using a transgenic line expressing plasma-membrane targeted GCaMP6s (Fig. 10 A; (Sheets et al., 2017)). In response to a 500 ms step deflection, calcium influx at the hair bundle was significantly reduced in dexamethasone-treated larvae relative to controls (Fig 10 B, B’; **P=0.0052; Unpaired t-test); an unexpected result given that uptake of the cationic dye FM1-43 was not reduced in dexamethasone-treated larvae (Fig. 5 A, B, E). By contrast, presynaptic calcium influx was not significantly different in dexamethasone-treated fish vs. controls (Fig. 10 C, D). Baseline GCaMP6sCAAX measurements at neuromast hair bundles and hair cell presynaptic regions were comparable between dexamethasone-treated and control animals (Fig. 10 E, F), indicating that calcium levels at rest were not altered in these two subcellular compartments. Overall, these results provide evidence that prolonged dexamethasone exposure does not affect hair cell calcium homeostasis but does negatively influence calcium flow through the hair bundle in response to stimulus.

**Figure 10.**
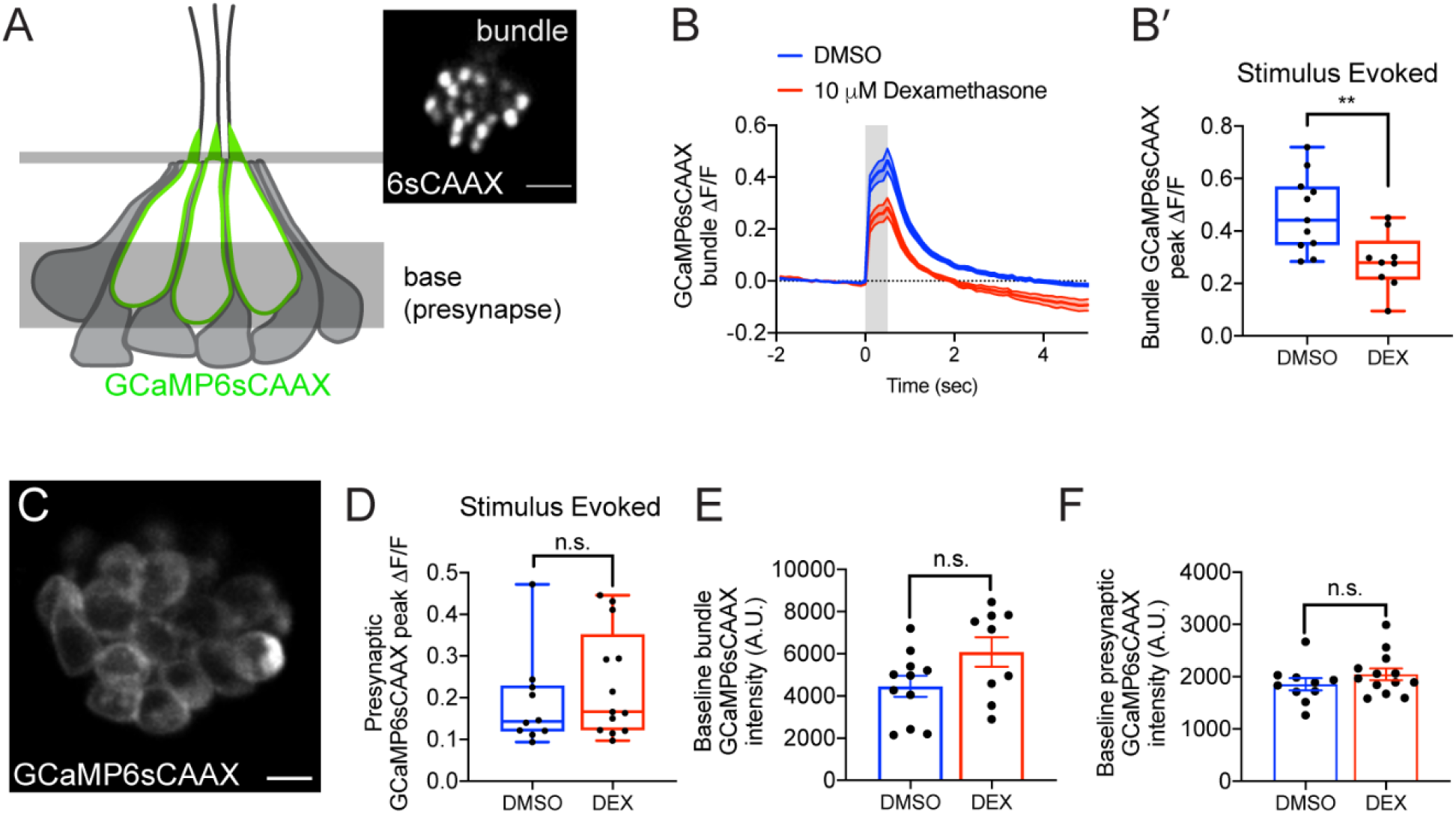
Dexamethasone exposure decreased evoked mechanosensation but not presynaptic calcium responses. (A) Schematic of a neuromast viewed from the side, and the imaging planes used to measure evoked calcium activity at the hair bundles and at the presynaptic region. Localization of GCaMP6sCAAX is shown in green. Inset shows an example of a top-down view of the hair bundle plane showing GCaMP6sCAAX fluorescence. (B-B’) Evoked hair bundle calcium signal with (red) or without (blue) 48-hour dexamethasone pretreatment. Average traces are shown in B, while dot plots of peak response of bundles per neuromast are shown in B’. (C) Top-down view of the basal plane showing the presynaptic membrane labeled with GCaMP6sCAAX. (D) Dot plots show that the peak evoked presynaptic calcium signal averaged per neuromast with (red) and without (blue) 48-hour dexamethasone pretreatment. (E-F) Baseline calcium intensity with (red) and without (blue) dexamethasone treatment in hair bundles (E) and presynapses (F) Shaded area above and below traces in panel B and error bars in E-F represent SEM. Box-and-whiskers plot were used in B’ and D to show median, min and max. An unpaired Welch’s unequal variance *t*-test was used to calculate the significance of differences in B’ and D. An unpaired *t*-test was used to calculate the significance of differences in E-F. n ≥ 9 neuromasts per treatment. **P=0.0052. Scale bar = 5 μm in A inset and C.

## Discussion

Dexamethasone is a potent synthetic glucocorticoid that influences diverse molecular pathways involved in cellular development, homeostasis, and repair. Using the zebrafish lateral line as a model for hair cell organs, we examined the influence of prolonged dexamethasone exposure on both afferent innervation of newly generated hair cells as well as hair cell homeostasis and function. Following 48 hours of dexamethasone exposure, we observed no inhibition of innervation or synapse formation in newly generated neuromast hair cells, indicating that dexamethasone did not inhibit regeneration of afferent synaptic contacts in the lateral-line system. However, further examination revealed prolonged dexamethasone exposure induced mitochondrial hyperpolarization in neuromast hair cells and supporting cells, suggesting disruption of mitochondrial homeostasis. In addition, dexamethasone exposure specifically reduced evoked calcium influx at the hair bundle but not the hair cell synapse in response to water-jet stimulus. Cumulatively, our data support that prolonged dexamethasone does not inhibit newly generated hair cell innervation or synapse formation, but it may negatively impact hair cell energy homeostasis and function.

### Dexamethasone did not inhibit lateral-line afferent nerve regrowth or hair cell synapse formation during regeneration

Exposure to dexamethasone significantly increased the total length of afferent neurites per regenerating neuromast (Fig 3 C-E). The addition in nerve fiber length corresponds to the increased number of hair cells (Fig. 3 F), suggesting that nerve fiber length was increased in regenerating neuromasts to accommodate a larger number of newly formed hair cells. Alternatively, it is possible that the additional neurites may be involved in facilitating hair cell generation. A recent study characterizing functional lateral-line neuromast regeneration following CuSO_4_ ablation has reported that afferent nerve terminals formed initial contacts with supporting cells during regeneration, suggesting that afferent nerve contacts may help facilitate transdifferentiation of supporting cells into hair cells (Hardy et al., 2021). However, while dexamethasone treatment caused regenerating neuromasts to add additional afferent dendrite length, no such increase was observed in unlesioned neuromasts (Fig. 3 A,B,E), suggesting that the generation of additional hair cells in unlesioned neuromasts is not facilitated by additional nerve fiber contacts. Given that dexamethasone treatment did not increase the number of afferent synapses per hair cell in unlesioned neuromasts (Fig 4 E; gray circles), we posit that afferent neurons innervate and form synapses with newly generated hair cells in unlesioned larval neuromasts without adding additional processes.

We also did not observe inhibition of macrophage activation in response to ototoxic damage following dexamethasone pretreatment (Fig 5). This result provides additional support to a previously reported observation that dexamethasone likely enhances neuromast regeneration via a mechanism other than immunosuppression (Namdaran et al., 2012). Instead, dexamethasone may be acting though glucocorticoid receptors to enhance division of a subset of hair cell precursors. Further studies will be needed explore the cellular pathways underlying dexamethasone-induced hair cell generation, and the mechanistic differences between afferent nerve innervation and synapse formation in regenerating vs. intact neuromasts.

### Prolonged dexamethasone exposure modulates mitochondrial membrane potential in neuromast cells and enhances neomycin hair cell toxicity

Glucocorticoid receptors reside in the cytosol of cells. When activated by a ligand such as cortisol or dexamethasone, they regulate gene expression by localizing to the cell nucleus and either directly binding to DNA or altering the activity of transcription factors or various kinases (Oakley & Cidlowski, 2013). Activated glucocorticoid receptors have also been shown to translocate into the mitochondria and regulate mitochondrial gene expression (Kokkinopoulou & Moutsatsou, 2021), and previous studies of mouse liver and rat distal colon have reported enhanced expression of mitochondrial genes involved in oxidative phosphorylation following dexamethasone treatment (Li et al., 2016; Rachamim et al., 1995). The glucocorticoid receptor gene *nr3c1* has been shown to be expressed in larval zebrafish neuromast hair cells and all populations of supporting cells (Lush et al., 2019). Speculatively, prolonged dexamethasone exposure may be modulating the expression of mitochondrial DNA encoded genes involved in oxidative phosphorylation, thereby altering mitochondrial metabolism in neuromast hair cells and supporting cells.

A recent study assessing the effects of cortisol exposure on hair cell sensitivity to neomycin further supports the hypothesis that activated glucocorticoid receptors regulates expression of genes that modulate mitochondrial metabolism (Hayward et al., 2019). Those authors reported that pretreatment of zebrafish larvae with cortisol for 24 hours sensitized neuromast hair cells to neomycin-induced hair cell death, which is comparable to what we observed in dexamethasone-treated fish (Fig. 9). Cortisol-mediated sensitization to neomycin was dependent on glucocorticoid receptor activation and could be blocked by inhibiting protein translation. It is worth noting that Hayward et al. did not report a similar sensitization to neomycin following 24 h dexamethasone exposure, but it is likely dexamethasone and cortisol are acting through a shared mechanism. The difference between their reported observations and our results is probably due to two factors. First, the concentration of neomycin used in Hayward et al. following dexamethasone was 25 μM, which was two-fold less than the dose at which we first observed significant effects ((Hayward et al., 2019); Fig 9 K). Also, the dexamethasone pretreatment doses (0.001-1 μM vs. 10 μM) and exposure time (24 vs. 48 hours) used by Haywood et al., (2019) were less than those used in the present study.

### Prolonged dexamethasone exposure reduces evoked calcium influx though mechanotransduction channels but does not affect hair cell calcium homeostasis

While we did observe hyperpolarization of mitochondrial membrane potential in dexamethasone-exposed neuromasts (Fig. 8), we did not observe significant changes in cytosolic or mitochondrial calcium homeostasis within hair cells relative to controls (Fig 7). By contrast, in response to water-jet stimuli, evoked calcium influx at the hair bundle was significantly reduced (Fig. 10 B), suggesting calcium influx through activated MET channels is reduced. This observed effect on calcium influx specifically at the hair bundle may be a consequence of altered mitochondrial metabolism. Mitochondria are important regulators of local intracellular calcium levels in sensory hair cells, and in addition to directly taking up calcium, mitochondria provide ATP to drive clearance of calcium though plasma membrane Ca^2+^-ATPases (PMCA1 and PMCA2), with PMCA2 shown to be densely localized in the hair cell stereocilia (Beurg et al., 2010). Zebrafish homologues of PMCA1 and PMCA2 are strongly expressed in mature neuromast hair cells of larval zebrafish (Lush et al., 2019). It is possible that altered mitochondrial metabolism may be influencing the activity of PMCA2 in dexamethasone-treated hair cells, thereby modifying PMCA2 regulation of calcium levels at the hair bundle. It is also conceivable that activated glucocorticoid receptors may be modulating expression of genes involved in mechanotransduction channel calcium permeability. Further experiments will be needed to determine the mechanism underlying dexamethasone’s specific effect on calcium influx through stimulated hair bundles.

In conclusion, our observations from zebrafish lateral-line neuromasts indicate that 48-hour dexamethasone treatment enhances regeneration and afferent reinnervation but alters mitochondrial homeostasis and evoked calcium influx through hair bundles. Dexamethasone is a steroid commonly used for treating hearing loss, and identifying strategies to retain high levels of dexamethasone for long periods within the inner ear is currently an active area of research. Considering our observations showing that prolonged exposure to dexamethasone had an adverse effect on hair cell mitochondrial metabolism and impacted hair bundle calcium influx, we believe future studies on the effects of long-term, localized dexamethasone treatment on mammalian cochlear function are indicated.

## Acknowledgments

This work was supported by the RNID (Project ID G85; L.S.), a National Institute on Deafness and Other Communication Disorders (NIDCD) Intramural Research Program Grant 1ZIADC000085-01 (K.S.K), and Extramural Research Project Grant R01DC006283 (M.E.W.).

